# Population genetics of Bull Trout (*Salvelinus confluentus*) in the Upper Athabasca river basin

**DOI:** 10.1101/743203

**Authors:** Emma K. T. Carroll, Steven M. Vamosi

**Affiliations:** Department of Biological Sciences, University of Calgary, 2500 University Drive NW, Calgary, Alberta, Canada T2N 1N4

**Keywords:** bull trout, Salvelinus, conservation, microsatellites, population genetics

## Abstract

Across its native range, Bull Trout (*Salvelinus confluentus*) extent and abundance are in decline due to historic overharvest and habitat degradation. Because Bull Trout are dependent on extensively connected, cold, clean headwater habitats, fragmentation from land use changes causes difficulty when determining the true extent and health of their populations, with Bull Trout of Alberta’s Eastern Slope region being no exception. Across this region, 431 Bull Trout from 20 sites were sampled from the Athabasca and Saskatchewan River basins and compared using 10 microsatellite loci to characterize within- and among-population genetic variation. The Saskatchewan and Athabasca River basins contained similar levels of heterozygosity but were differentiated from one another. Within the Athabasca River basin, five genetically differentiated clusters were found. Additionally, no isolation-by-distance pattern was observed between these sites. These results suggest these populations have ample genetic diversity, but genetic differentiation should be considered when deciding whether and how to alter connectivity between populations.

## Introduction

Quantification and incorporation of population genetic information can provide valuable guidance when undertaking conservation efforts to protect populations of a declining species (Epifanio et al. 2003). When populations contain sufficient genetic diversity, natural selection can act on beneficial alleles to facilitate local adaption (Frankham 2005; Tiffin & Ross-Ibarra 2014). Especially if habitat fragmentation is occurring, functional connectivity and gene flow between populations typically decrease, constricting the species range and increasing isolation between populations (Pierce *et al*. 2013). These demographic changes have genetic consequences, from population differentiation to genetic bottleneck effects and inbreeding, especially at low population densities (Neraas & Spruell 2001). Loss of genetic variation, whether allelic richness or heterozygosity, exposes the population to inbreeding effect and genetic drift, both of which further decrease genetic diversity and heterozygosity through allele fixation (Frankham 2005). If allele fixation occurs, the potential for expression of deleterious recessive alleles increases, with subsequent decrease in fitness related traits in that population (Ruiz-López *et al*. 2012). In the absence of genetic rescue via gene flow from migrants, these negative genetic consequences can lead to population extirpation (Höglund 2009).

Bull Trout (*Salvelinus confluentus*, Suckley 1895) is an apex predator native to northwestern North America. Across this vast range, spanning different watersheds, ecoregions and management areas, the species is declining in population size and range extent due to overharvest, reduced habitat connectivity, and habitat degradation (Taylor *et al*. 1999). Owing to Bull Trout’s important ecological role and popularity as a recreational fishery, we have additional incentive to understand and monitor this sensitive, cold-water dwelling species (Warnock, Blackburn & Rasmussen 2011). Due to rarity and seasonal habitat use, population genetic approaches to studying Bull Trout have provided a better understanding of how the species is responding to stressors (Taylor *et al*. 1999; Costello *et al*. 2003; Ardren *et al*. 2011), primarily in the southern extent of their range. This research has shown that, in common with other Salmonids, a single Bull Trout population can contain multiple life history forms and migratory behaviours that constitute a single reproductive unit (Homel *et al*. 2008). Resident forms inhabit natal headwater streams, where a main conservation concern is the extirpating potential due to industrial caused alterations to habitat (Ripley *et al*. 2005). Additional natural sources of headwater habitat changes include stochastic events including debris flows and fire, both common in headwater stream (Rieman & Mcintyre 1995). Migratory forms travel downstream to mainstem rivers (fluvial form), lakes (adfluvial) or the ocean (anadromous) where they access their adult foraging habitat (Taylor *et al*. 1999). Bull Trout populations require extensively connected freshwater passageways to allow individuals of each migratory strategy access to spawning and foraging habitats (DeHaan *et al*. 2011). Consequently, the range of a single individual can be up to 500 km across highly heterogeneous waterways (Pillipow & Williamson 2003) while another in the same population will remain in the natal stream their entire life (Ardren *et al*. 2011). Due to spawning site fidelity, Bull Trout population genetics typically exhibit hierarchical population structure (Evanno *et al*. 2005), whereby high levels of genetic differentiation between populations causes population sub-structuring within a river basin (Whiteley *et al*. 2004).

Genetic tools can help us describe, understand, and monitor how Bull Trout populations are coping with the biological and physical alterations to their habitat on local (Ardren *et al*. 2011) and range-wide (Taylor *et al*. 1999) scales. Rare gene flow between these spawning streams typically results in a positive, albeit weak correlation between distance and genetic similarity, although populations typically contain unique allelic variants (Warnock *et al*. 2010). Because threatened populations typically contain less genetic heterozygosity than species of non-concern, it is crucial to detect these trends of declining genetic diversity while correction measures are still viable (Spielman *et al*. 2004).

At present, there are gaps in our understanding of Bull Trout in the eastern portion of its range, east of the continental divide, specifically the Athabasca River basin (Ripley *et al*. 2005). In this region, two of five of Canada’s Designated Units (DUs) persist in allopatry: the well-studied Saskatchewan-Nelson River population (DU4) to the south and the understudied Western Arctic population (DU2) to the north (COSEWIC 2012). Currently, there is relatively little data on the genetic diversity or genetic relationships within and among these Bull Trout populations, especially in the headwaters region (Taylor *et al*. 1999). Evaluating the status of a species’ genetic diversity through population genetic techniques enables us to detect population trends, such as inbreeding depression (Restoux *et al*. 2012), hybridization (Rhymer & Simberloff 1996), and population isolation by a cryptic barrier (Bergek & Björklund 2007). With reductions to population numbers and to the overall range of Bull Trout, this gap in our knowledge of their genetics negatively affects our ability to identify vulnerable populations and adequately protect them.

Here, we assess the population genetic structure of Bull Trout, a globally recognized species as risk (COSEWIC 2012), in the upper reaches of the Athabasca River basin. We used neutral markers to genetically identify putative populations and characterize genetic differentiation within and among populations and among river basins in Alberta. Our specific objectives were to: (1) identify Bull Trout populations in the Athabasca River basin, and (2) characterize within- and among-population genetic differentiation for three major river basins (Athabasca, North Saskatchewan, and Bow), which have evolved from the same genetic lineage (COSEWIC 2012). Within the Athabasca River Basin, we expect that Bull Trout will follow the typical hierarchical population structuring seen in other river basins where intra-population genetic differentiation is low, inter-population genetic differentiation is high, and isolation by distance is weak. By measuring genetic diversity and differentiation, these data will inform management decisions by establishing a baseline dataset. With this dataset, we will have the ability to identify genetically depauperate populations and extreme differentiation, which may suggest barriers to migration and gene flow. Overall, we provide insight into how cryptic diversity in Bull Trout may inform local management and conservation strategies.

## Methods

### Study location and sample collection

We sampled 14 sites to characterize the genetic diversity of the Bull Trout populations within the Athabasca River Basin (Western Arctic), with six sites from the North Saskatchewan and Bow River Basins (Saskatchewan-Nelson) sampled for comparison (Figure 1). Sites were chosen based on accessibility and known presence of Bull Trout. Both fluvial and adfluvial sites were sampled, thus fish of different life history strategies are combined in subsequent analyses. Bull Trout ranged in fork length size from 51mm (Berland River) - 610 mm (Athabasca River). A total of 431 individuals were captured from 2007-2015 and were sampled at 20 locations (Table 1). Fish were caught from adfluvial sites by angling and by either backpack or boat electrofishing at fluvial sites. Samples were collected from each Bull Trout via non-lethal adipose fin tissue collection. Upon capture, fork length measurements were taken of each individual and adipose fin tissue samples were clipped and transferred into 95% EtOH (Ethanol) at 4°C. Samples obtained from W. Hughson, S. Herman, M. Sullivan and W. Warnock (Table 1) were collected using similar methods, although the latter three collectors preserved the sample by drying the adipose fin tissue and storing them in separate envelopes in a cool, dry location. All samples were collected within eight years, a time span comparable to a single Bull Trout generation within the region. Fish collection was completed under approved fish research licences (Parks Canada: 18570 and Alberta Environment and Parks: 15-2028) and animal care protocol from Life and Environmental Sciences Animal Resource Centre at the University of Calgary (AC15-0033).

**Figure 1:**
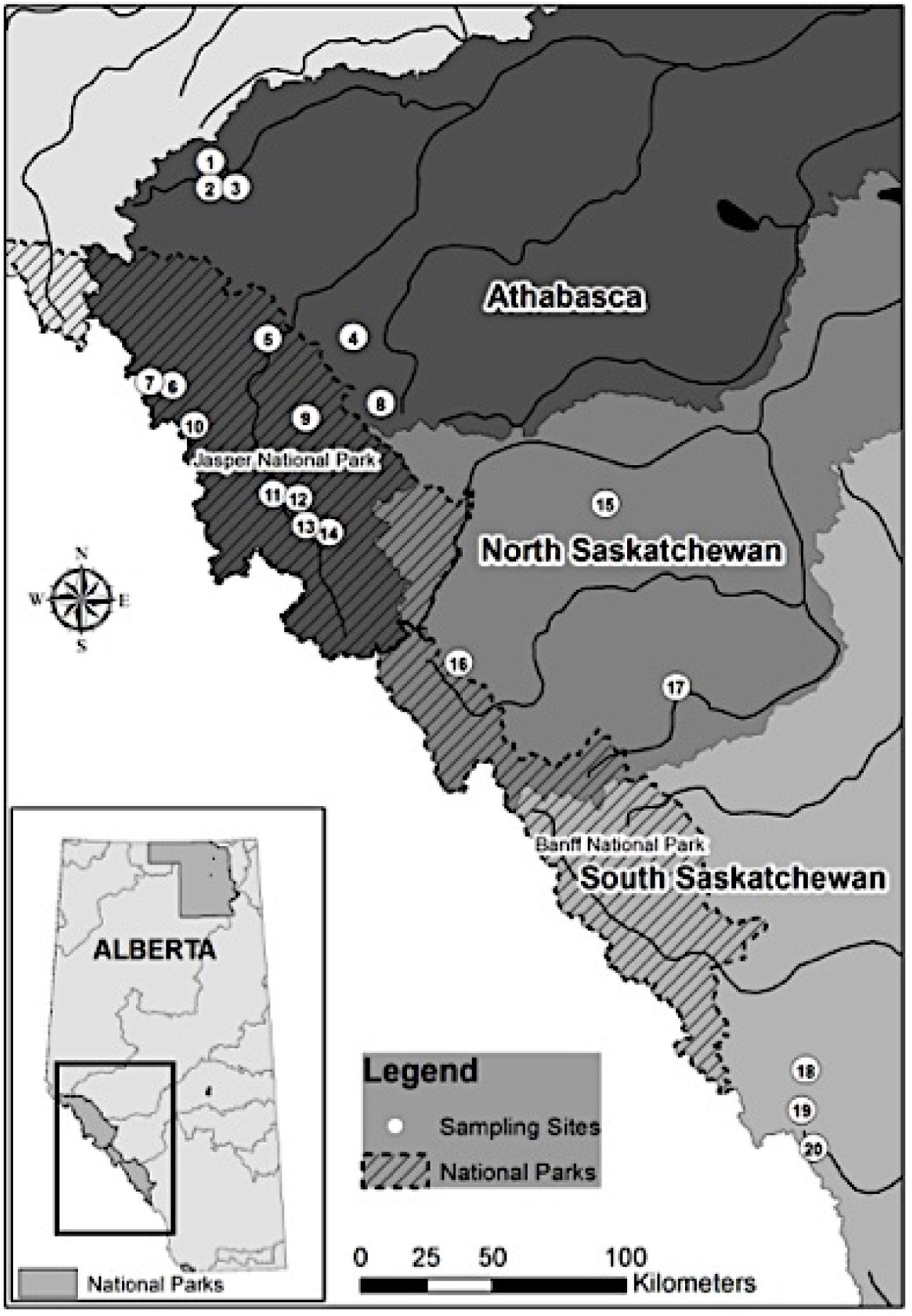
Sample sites for Bull Trout in Alberta, Canada in the Athabasca, North Saskatchewan and Bow watersheds. Colours denote different watersheds, numbers represent sampling sites (Table 1).

**Table 1:** Collection data summary. Sampling year are denoted as follows: + = 2007, ⊙= 2010, ❖ = 2011, ▪ = 2015). Site ID and abbreviations (Abr) are used in Figure 1. BKTR denotes whether Brook Trout (BKTR) was also found in the sampled areas.

| ID | Watershed | Site | Abr | Latitude | Longitude | *n* | Collector | BKTR |
| --- | --- | --- | --- | --- | --- | --- | --- | --- |
| 1 | Athabasca | Berland River | BR | 53.7677 | -118.3523 | 25 | Carroll■ | Yes |
| 2 | Athabasca | Moon Creek | MC | 53.7050 | -118.3502 | 25 | Carroll■ | No |
| 3 | Athabasca | Little Berland River | LBR | 53.6798 | -118.2373 | 25 | Carroll■ | No |
| 4 | Athabasca | Gregg River | GR | 53.2017 | -117.5007 | 17 | Carroll■ | Yes |
| 5 | Athabasca | Athabasca River | AR | 53.1861 | -117.9821 | 25 | Hughson■ | Yes |
| 6 | Athabasca | Miette Lake | ML | 53.0223 | -118.5585 | 25 | Sullivan+ | No |
| 7 | Athabasca | Unnamed Lake | UL | 53.0182 | -118.6316 | 10 | Sullivan+ | No |
| 8 | Athabasca | McLeod River | MR | 52.9847 | -117.335 | 25 | Carroll■ | Yes |
| 9 | Athabasca | Jacques Lake | JL | 52.9274 | -117.7522 | 25 | Carroll■ | No |
| 10 | Athabasca | Derr Creek | DC | 52.8853 | -118.3761 | 1 | Carroll■ | Yes |
| 11 | Athabasca | Kerkeslin Creek | KC | 52.6799 | -117.8709 | 25 | Carroll■ | Yes |
| 12 | Athabasca | Kerkeslin Lake | KL | 52.6573 | -117.7875 | 25 | Carroll■ | No |
| 13 | Athabasca | Ranger Creek | RC | 52.5733 | -117.7160 | 15 | Carroll■ | Yes |
| 14 | Athabasca | Osprey Lake | OL | 52.5592 | -117.6697 | 25 | Carroll■ | No |
| 15 | North Saskatchewan | Colt Creek | CC | 52.6667 | -116.0689 | 25 | Herman■ | Yes |
| 16 | North Saskatchewan | Pinto Lake | PL | 52.1248 | -116.8637 | 25 | Herman■ | No |
| 17 | North Saskatchewan | Elk Creek | EC | 52.0586 | -115.6663 | 25 | Herman■ | Yes |
| 18 | Bow | Little Elbow River | LER | 50.7796 | -114.9537 | 25 | Warnock❖ | Yes |
| 19 | Bow | Elbow River | ER | 50.6748 | -114.9757 | 13 | Warnock❖ | Yes |
| 20 | Bow | Storm Creek | SC | 50.5187 | -114.9126 | 25 | Warnock⊙ | Yes |

### DNA sequence amplification

Nuclear DNA (Deoxyribonucleic Acid) was extracted from EtOH-preserved adipose fin tissue using a standard proteinase-K phenol-chloroform protocol (Sambrook *et al*. 1989) and stored in ddH2O at 4°C. DNA quantity (μg/μl) was determined using a Nanodrop 1000 Spectrophotometer V3.7 (Thermo Fisher Scientific). DNA quantity was measured and sample concentrations were adjusted to 50 ng/μl, the desired concentration for the PCR (Polymerase Chain Reaction) protocol (New England Biolabs Inc.).

During PCR, DNA regions, excised by forward fluorescently labeled oligonucleotide primers for 10 microsatellite loci (Omm1128, Sco102, Sco105, Sco106, Sco109, Sco215, Sco216, Sco220, Sfo18, and Smm22), were multiplexed in four optimized groups (Sup. Table1). These loci were chosen based on the degree of polymorphism, ability to detect Bull Trout × Brook Trout hybrids, and resolution to detect population structure of the samples (Warnock *et al*. 2010). Desired fragments were amplified using a C1000 Touch Thermal Cycler (Bio-Rad, Inc., Hercules, CA).

PCR products were run on a 1.5% w/v agarose gel using a standard gel electrophoresis protocol and visualized using DigiDOC IT electrophoresis gel imager (UVP Inc., Upland, CA). Samples showing clearly defined bands representing the DNA fragment PCR products of the specified microsatellite loci were then analyzed. Fragment lengths were determined using a standard protocol for microsatellite fragment analysis with an ABI 3500XL Capillary Electrophoresis Genetic Analyzer (Applied Biosystems, Inc.) and scaled against GeneScan-500LIZ size standard (Applied Biosystems, Inc.). For each sample, electropherograms were produced, which were scored using GENEMAPPER v.4.1 (Applied Biosystems, Inc.).

### Genetic data analysis

To quantify human scoring error and machine processing error, we followed recommendations set out by Guichoux *et al*. (2011). To quantify human scoring error, original raw electropherograms of 500 of 4310 (11%) images were rescored and the resulting genotypes were compared to the initially scored electropherogram. Clear electropherograms and strict scoring criteria (Guichoux *et al*. 2011) contributed to no human scoring differences between the two scoring rounds. Replicates of 711 electropherograms of the total 4310 (16%) were run and scored, revealing a 1.7% machine error rate, which is comparable to other studies using tissue for DNA extractions (Morin *et al*. 2007). These loci were disregarded, but an equal error is still presumed to be present in the unchecked portion of the data set.

Raw genotype scores were assessed by MICROCHECKER v.2.2.3 (Van Oosterhout *et al*. 2004) to uncover genotyping errors and presence of null alleles that did not amplify during PCR. Because null alleles were present in no more than four of the 10, all samples were used for further analyses loci (Van Oosterhout *et al*. 2004).

### Hierarchical population structure

Bull Trout tend to exhibit hierarchical population structure (Evanno *et al*. 2005; Warnock 2008), whereby high levels of genetic differentiation between populations causes population sub-structuring (Whiteley *et al*. 2004). Therefore, among river basin relationships and within and among sub-basin relationships were evaluated using the Bayesian clustering method in STRUCTURE v.2.3.4 (Pritchard et al. 2000). All analyses in STRUCTURE utilized the correlated allele frequency model and flexibility in linkage disequilibrium parameters to allow the complexities of natural systems to be included into the STRUCTURE estimates (Evanno *et al*. 2005; Vähä *et al*. 2007). These parameters allow allele frequencies to be correlated between populations and are able to accurately assign individuals to closely related populations due to admixture or recent common ancestry (Falush *et al*. 2003). This scenario is biologically plausible in this study system due to the potential for mainstem river sampling sites to contain migratory adults from different populations that could be classified as a single population.

Including all sites within the Athabasca and Saskatchewan watersheds, 10 independent STRUCTURE runs (Evanno *et al*. 2005) performed at *K*-values of 1-20 were performed to confirm genetic differentiation of Bull Trout between major river basins, the Athabasca and Saskatchewan. Subsequent STRUCTURE analyses were performed on Bull Trout captured within the Athabasca River basin to determine sub-basin structure using 10 independent runs at *K*-values of 1-14. We determined the ranges of *K*-values based on the potential of all samples to be part of a panmictic system (*K*_MIN_ =1) and the assumption that STRUCTURE detects the major structure, which is uncommonly greater than the total number of sampling sites (*K*_MAX_ = *N*_sites_) (Pritchard *et al*. 2000). For all *K*-values in STRUCTURE, 10 independent replicates were run. Initial STRUCTURE runs were conducted at 100,000 burn-in lengths and 100,000 Markov chain Monte Carlo (MCMC) iterations (Warnock 2008) to reveal coarse, large-scale structure among the water basins while subsequent sub-watershed STRUCTURE runs were conducted at 500,000 burn-in lengths and 500,000 MCMC iterations to give more resolution to the localized population structure within watersheds (Warnock 2008).

To determine the optimal value of *K,* the simulation summary results of the 10 independent runs for each value of *K* were evaluated with Structure Harvester v0.6.94 (Earl & vonHoldt 2012). If multiple unique STRUCTURE plots were created at each *K*, CLUMPAK (Kopelman *et al*. 2015) was used to find the optimal alignment of runs at a given value of *K* using CLUMPP (Jakobsson and Rosenberg 2007) and plotted using DISTRUCT (Rosenberg 2004). Using the recommendations of Pritchard *et al*. (2000) and Evanno *et al*. (2005), the model in which the lowest value of *K* that encompassed the majority of the structure while also having the highest rate of change in the log probability of the data (Δ*K*) was deemed the most likely correct *K* value. The hierarchical partitioning of genetic variation among river basins and drainages using Analysis of Molecular Variance (AMOVA) in GenAlEx version 6.5 (Peakall & Smouse 2006).

### Population genetics

For all Bull Trout sampled across all sites containing >10 individuals, departures from Hardy-Weinberg Equilibrium (HWE), heterozygote excess and deficiency, and Linkage Disequilibrium (LD) between pairs of loci were determined using Markov-chain methods and the following parameters in Genepop version 4.2 (Rousset 2008): MCP – 100,000 dememorizations, 5,000 iterations (Warnock 2008). Both HWE and LD tests levels of significance were adjusted using nonsequential Bonferroni corrections (Rice 1989). Genetic diversity was calculated across all sites with genetic diversity and private alleles calculated in GenAlEx version 6.5 (Peakall & Smouse 2006) and allelic richness in FSTAT version 2.9 (Goudet 2001). Genetic divergence *F*_ST_ and Inbreeding *F*_IS_ (Weir and Cockerham 1984) was estimated in FSTAT 2.9 (Goudet 2001). *F_ST_* and *F*_IS_ significance levels were adjusted using nonsequential Bonferroni correction (Rice 1989).

Isolation by distance was tested for in the Athabasca River basin using the Mantel Test to determine the relationship between pairwise genetic distance (*F_ST_*) and geographic distance (km) (Jensen *et al*. 2005) with 1000 randomizations in each scenario. Geographic distances were obtained using GoogleEarth (v. 4.2.1, Google Inc.).

## Results

### Microsatellite loci

The degree of polymorphism displayed by the microsatellite loci of the 431 Bull Trout ranged from 7 (Sfo18) to 51 (Sco109). Genotypic frequencies differed from HWE in a number of locations and alleles. Heterozygote excess was rare and found only in Little Elbow River (Sco102) and Storm Creek (Omm1128). Evidence for heterozygote deficiency was found in the populations from: Colt Creek (Sco102, Sfo18), Elk Creek (Sfo18), Kerkeslin Lake (Sfo18, Sco220), Miette Lake (Sfo18), Moon Creek (Sco102, Sfo18), and Storm Creek (Sco109). One pair of loci (JL; Sco115xSco220) was observed at one of 450 total locus pairs across sites to be in linkage disequilibrium after Bonferroni correction, with no linkage trends between any two loci across all sites and loci. Infrequent evidence for null alleles was seen throughout the dataset. (Sup. Table 2, Sup. Table 3; online supplemental material)

The two loci used to differentiate Bull Trout from Brook Trout and their hybrids, Sco216 and Sfo18, all scored alleles outside the range indicative of Brook Trout DNA, 172-192bp and 145-165bp respectively (Popowich *et al*. 2011) corroborating that all tissues are from genetically pure Bull Trout.

### Population genetics

Genetic diversity was not significantly different between the Athabasca River basin (*H_E_* = 0.585) and Saskatchewan River basin (*H_E_*=0.567; Welch’s *t*-test, *t*=0.5002, *df*=11.149, *P*=0.627). In the Athabasca River basin, expected heterozygosity values ranged from 0.473 (Gregg River) to 0.736 (Miette Lake). Allelic richness values averaged 5.52 and ranged from 2.8 (Unnamed Lake) to 8.6 (Athabasca River). Inbreeding coefficient (F_IS_) ranged from −0.122 (Ranger Creek) to 0.152 (Moon Creek). Of the 41 private alleles detected in this study, the Athabasca contained 20, with values ranging from zero (Little Berland River and Ranger Creek) to four (Kerkeslin Creek and Unnamed Lake). Sites within the Saskatchewan River basin had similar expected heterozygosity, ranging from 0.468 (Elbow River) to 0.690 (Colt Creek). Allelic richness averaged 6.5 and ranged from 3.7 (Elbow River) to 8.1 (Colt Creek). The Saskatchewan River basin contained the remaining 21 private alleles detected, with values ranging from zero (Colt Creek, Storm Creek, and Elbow River) to nine (Little Elbow River).

### Hierarchical population structure

#### Among-river basin structure (broad scale)

Genetic differentiation was detected in the Athabasca River basin and Saskatchewan River basin samples although no differences in levels of heterozygosity were observed between the river basins. The mean *F_ST_* value within the Athabasca River basin was 0.25, ranging from 0.05 (between Athabasca River and Kerkeslin Creek) to 0.49 (between McLeod River and Kerkeslin Lake. The mean *F_ST_* in the Saskatchewan River basin was similar (0.24) to the Athabasca river basin samples, with *F_ST_* values ranging from 0.15 (between Storm Creek and Little Elbow River) to 0.33 (between Elbow River and Pinto Lake) (Table 3). Using the clustering method, STRUCTURE analyses revealed a strong signal at *K*=2 (Δ*K* = 13.36, Figure 2), representing the coarsest level of structure within the area sampled. These two distinct clusters aligned with the geographical split of major river basins: one cluster contained all samples from the Athabasca River basin (Western Arctic DU) (Sites 1-14), whereas the second cluster encompassed all samples from the Saskatchewan River basin (Saskatchewan-Nelson DU; North Saskatchewan: Sites 15-17; Bow: Sites 18-20; Table 1, Figure 3). Although clustered based on river basin, the majority of regional variation was explained when sampling sites were grouped based on their location within the Athabasca River basin and the Saskatchewan River basin (Table 2). Additionally, an Analysis of Molecular Variance (AMOVA) showed that across all measured regions, both among and within major river basins, the greatest amount of genetic variation was explained by population level groupings (Table 2).

**Figure 2:**
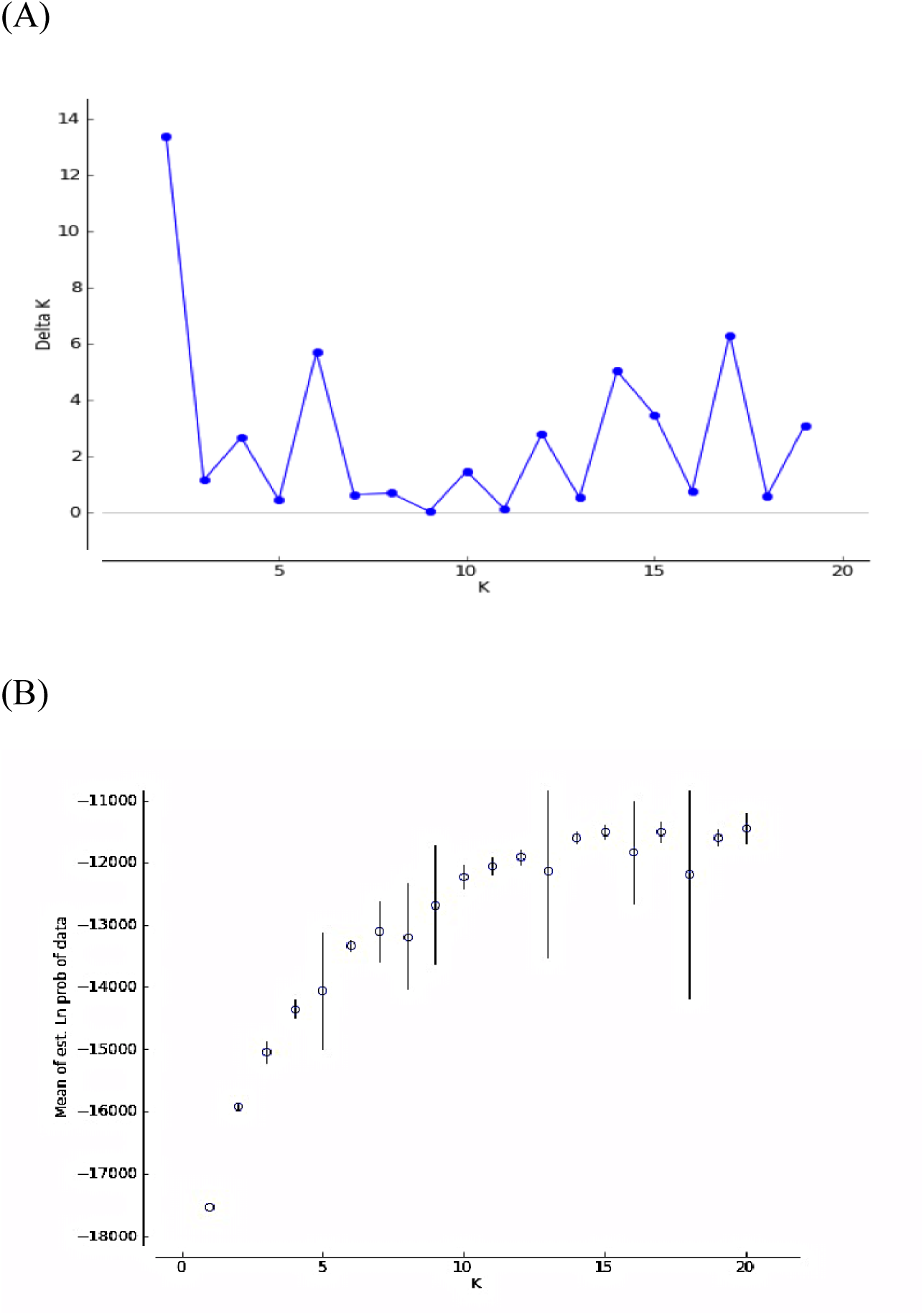
(A) STRUCTURE results of a distribution of *DeltaK* (the second order rate of change in the log probability of the data; based on Evanno *et al*. 2005) for all 20 sites sampled across the Athabasca and Saskatchewan River basins. *K* represents the number of clusters detected in the data set. (B) Posterior probability (LnP(D)) per cluster (*K*) as recommeded by Pritchard et al. (2000) with standard error bars for all 20 sites sampled across the Athabsca and Saskatchewan River basins

**Figure 3:**
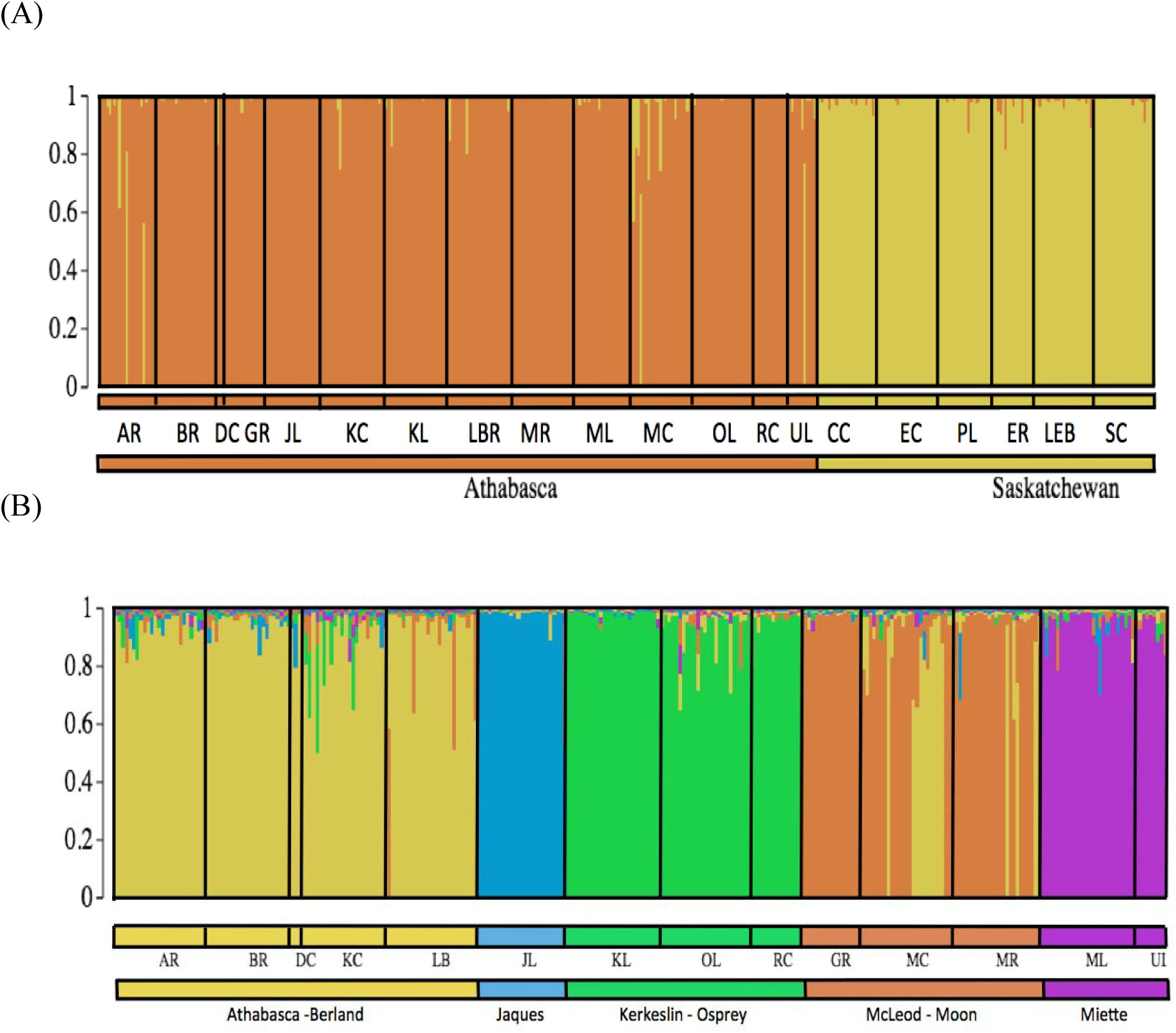
STRUCTURE result of admixture plots for Bull Trout sampled in (A) the Athabasca and Saskatchewan River basins and subbasin structure within the Athabasca River basin only (B). Admixture plots showing individual membership to *K* clusters for *K* values of 2 (A) and 5 (B) based on genotypes. Unique clusters are represented by colour. Each vertical line represents an individual. See Table 1 for sampling site codes.

**Table 2:**
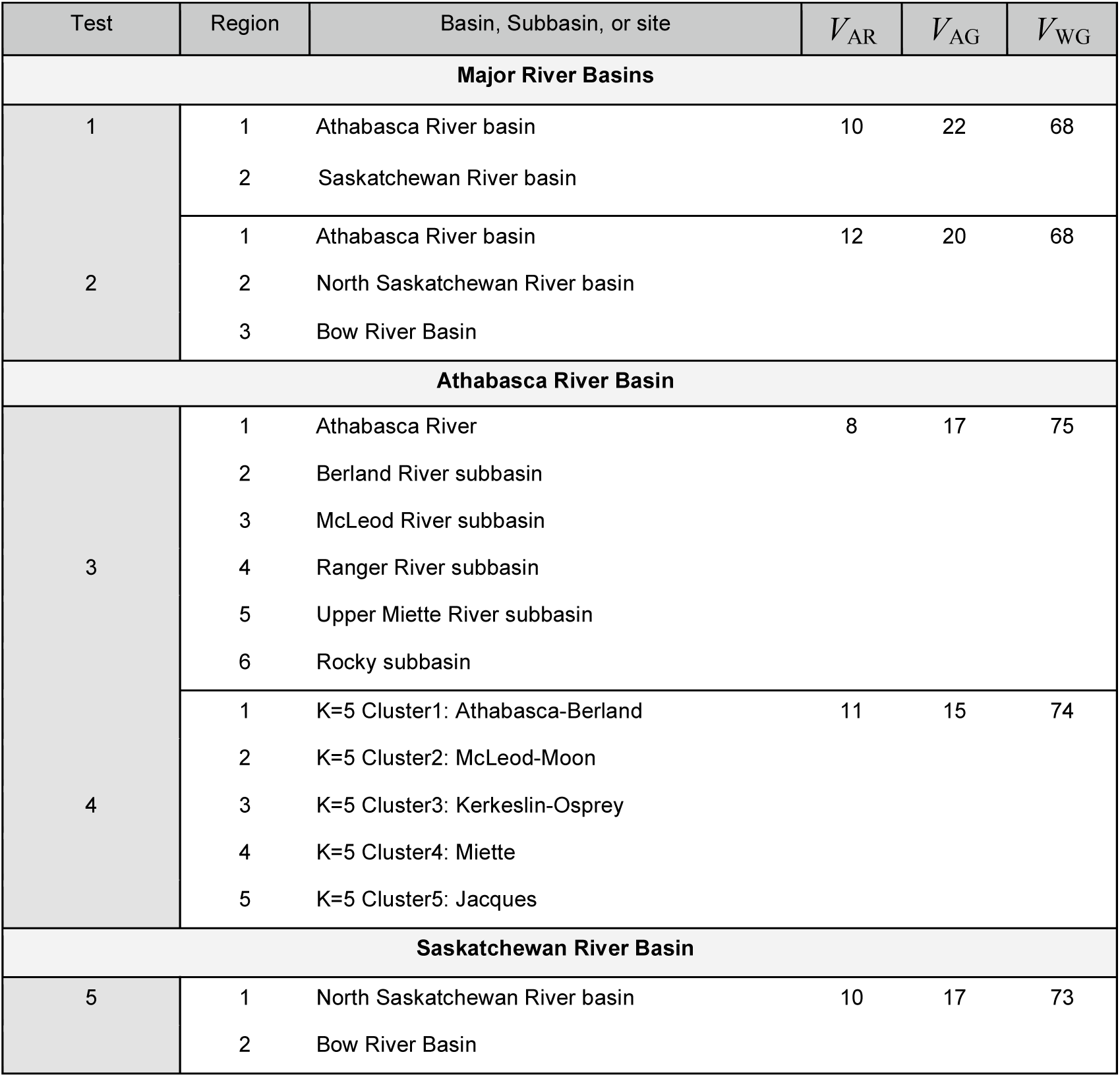
Bull Trout genetic variance subdivided by Analysis of Molecular Variance (AMOVA; Excoffier and Lischer 2010) explained among regions (*V*_AR_), among groups (*V*_AG_), and within groups (*V*_WG_).

**Table 3:** Pairwise genetic differentiation index *F_ST_* estimates (Weir and Cockerham 1984) among sample sites. All sites pairs are significantly different from panmixia (P<0.0003). See population codes in Table 1.

| Site Codes | AR | BR | CC | EC | ER | GR | JL | KC | KL | LER | LBR | MR | ML | MC | OL | PL | RC | SC | UL |
| --- | --- | --- | --- | --- | --- | --- | --- | --- | --- | --- | --- | --- | --- | --- | --- | --- | --- | --- | --- |
| AR | - |  |  |  |  |  |  |  |  |  |  |  |  |  |  |  |  |  |  |
| BR | 0.06 | - |  |  |  |  |  |  |  |  |  |  |  |  |  |  |  |  |  |
| CC | 0.20 | 0.23 | - |  |  |  |  |  |  |  |  |  |  |  |  |  |  |  |  |
| EC | 0.27 | 0.31 | 0.18 | - |  |  |  |  |  |  |  |  |  |  |  |  |  |  |  |
| ER | 0.30 | 0.32 | 0.26 | 0.33 | - |  |  |  |  |  |  |  |  |  |  |  |  |  |  |
| GR | 0.17 | 0.16 | 0.25 | 0.38 | 0.42 | - |  |  |  |  |  |  |  |  |  |  |  |  |  |
| JL | 0.23 | 0.21 | 0.31 | 0.37 | 0.43 | 0.27 | - |  |  |  |  |  |  |  |  |  |  |  |  |
| KC | 0.06 | 0.07 | 0.19 | 0.26 | 0.29 | 0.19 | 0.22 | - |  |  |  |  |  |  |  |  |  |  |  |
| KL | 0.36 | 0.39 | 0.33 | 0.33 | 0.48 | 0.46 | 0.45 | 0.30 | - |  |  |  |  |  |  |  |  |  |  |
| LEB | 0.27 | 0.31 | 0.25 | 0.29 | 0.17 | 0.37 | 0.39 | 0.27 | 0.45 | - |  |  |  |  |  |  |  |  |  |
| LBR | 0.10 | 0.06 | 0.26 | 0.36 | 0.38 | 0.19 | 0.30 | 0.13 | 0.45 | 0.34 | - |  |  |  |  |  |  |  |  |
| MR | 0.13 | 0.12 | 0.30 | 0.39 | 0.44 | 0.21 | 0.33 | 0.16 | 0.49 | 0.38 | 0.11 | - |  |  |  |  |  |  |  |
| ML | 0.27 | 0.31 | 0.28 | 0.32 | 0.39 | 0.35 | 0.42 | 0.27 | 0.47 | 0.31 | 0.30 | 0.32 | - |  |  |  |  |  |  |
| MC | 0.09 | 0.08 | 0.21 | 0.30 | 0.29 | 0.13 | 0.26 | 0.12 | 0.41 | 0.25 | 0.08 | 0.14 | 0.27 | - |  |  |  |  |  |
| OL | 0.15 | 0.18 | 0.25 | 0.35 | 0.38 | 0.23 | 0.32 | 0.13 | 0.35 | 0.33 | 0.19 | 0.23 | 0.31 | 0.17 | - |  |  |  |  |
| PL | 0.26 | 0.28 | 0.22 | 0.24 | 0.33 | 0.35 | 0.33 | 0.24 | 0.42 | 0.31 | 0.35 | 0.37 | 0.34 | 0.29 | 0.34 | - |  |  |  |
| RC | 0.24 | 0.22 | 0.29 | 0.36 | 0.40 | 0.30 | 0.35 | 0.18 | 0.41 | 0.38 | 0.28 | 0.34 | 0.40 | 0.20 | 0.17 | 0.35 | - |  |  |
| SC | 0.27 | 0.31 | 0.23 | 0.25 | 0.18 | 0.36 | 0.37 | 0.26 | 0.38 | 0.15 | 0.37 | 0.41 | 0.35 | 0.28 | 0.34 | 0.28 | 0.35 | - |  |
| UL | 0.27 | 0.30 | 0.24 | 0.32 | 0.34 | 0.36 | 0.44 | 0.26 | 0.48 | 0.30 | 0.32 | 0.37 | 0.24 | 0.25 | 0.31 | 0.31 | 0.36 | 0.34 | - |

#### Within-river basin structure (fine scale)

Within the Athabasca River basin, further genetic differentiation revealed additional levels of structuring. Cluster-based analyses in STRUCTURE revealed five sub-basin archipelagos, each containing samples from one or more drainages within it (Figure 2). From the 14 sampling sites in this watershed, the strongest signal occurred at *K*=5 (Δ*K* =196.96; Figure 3, Figure 4). These clusters aligned reasonably well with geographical drainages present in the water basin (Table 1), clustering sites from the Rocky drainage (Jacques Lake) and Miette drainage (Miette and Unnamed Lake) to their respective drainages. This supports local supposition of impassable barriers between the sampled areas relative to the mainstem Athabasca River (Ward Hughson, Pers. Comm.) The three other clusters each contained fish sampled from sites spanning larger areas and different drainages. The three mixed drainage clusters were comprised of: (1) the majority of sites from the Berland drainage and the main stem Athabasca drainage (Athabasca-Berland), (2) the upper Kerkeslin drainage site with all the Ranger drainage sites (Kerkeslin-Osprey), and (3) all McLeod drainage sites with an upper Berland drainage site (McLeod-Moon). We note that *K*=5 clusters accounted for more genetic variation than grouping sites by drainages (Table 2). Regardless of regional grouping, the majority of the genetic variation was explained within sampling sites.

**Figure 4:**
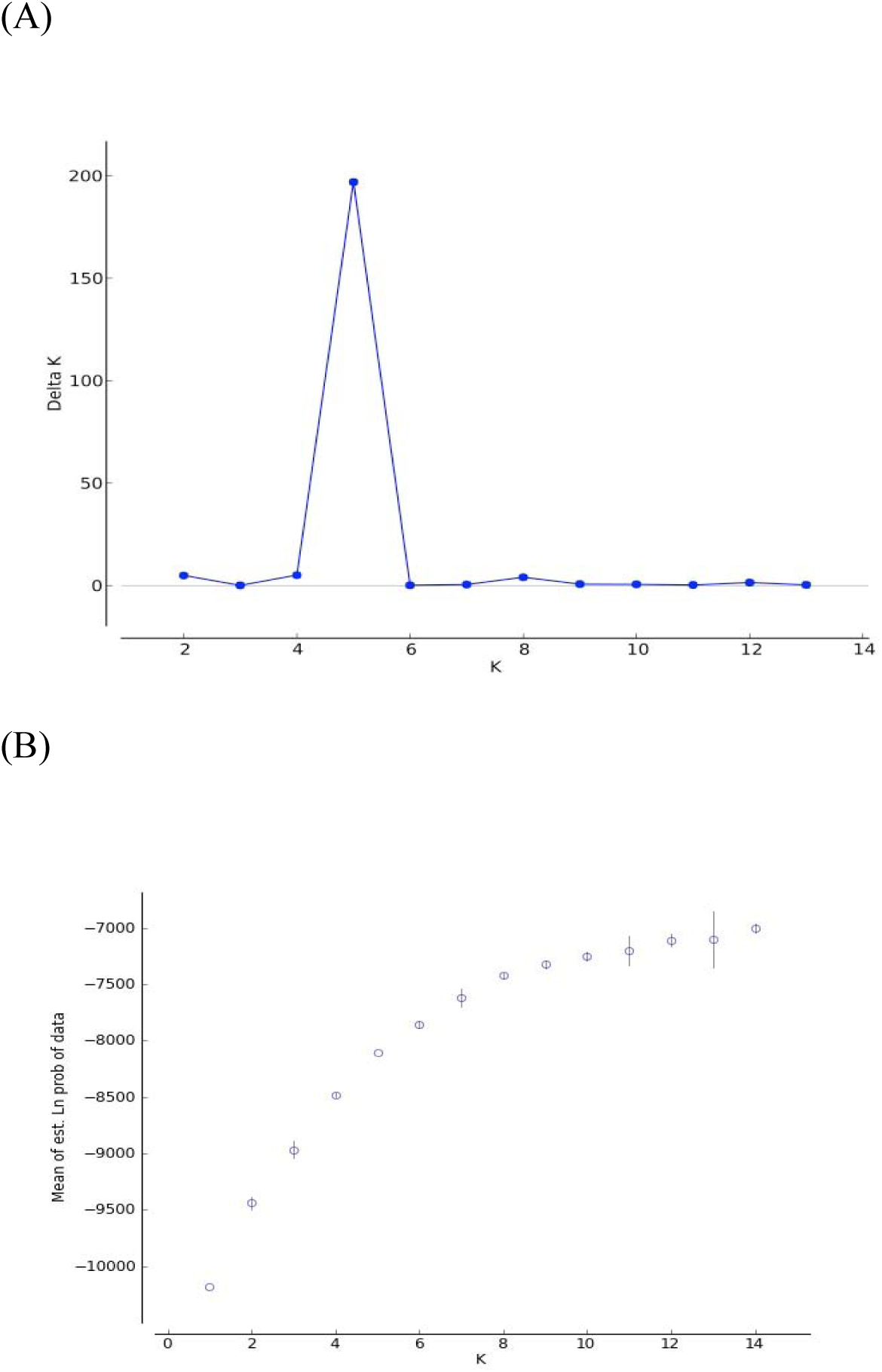
(A) STRUCTURE results of a distribution of *DeltaK* (the second order rate of change in the log probability of the data; based on Evanno *et al*. 2005) for only the 14 sites sampled across the Athabasca River basins. *K* represents the number of clusters detected in the data set. (B) Posterior probability (LnP(D)) per cluster (*K*) as recommeded by Pritchard et al. (2000) with standard error bars for all 14 sites sampled across the Athabsca River Basin

### Isolation by Distance

In the Athabasca River basin, pairwise *F_ST_* values were variable at all geographic distances (Figure 5). Mantel tests performed on Athabasca River basin Bull Trout revealed no significant isolation-by-distance pattern between genetic distance and geographic distance using either waterway distance (Figure 5; *Z*=45.78, *r*=0.079, *P*=0.29, *R*^2^=6.2e-3) or linear distance (*Z*=36.69, *r*=0.072, *P*=0.19, *R*^2^=5.1e-3).

**Figure 5:**
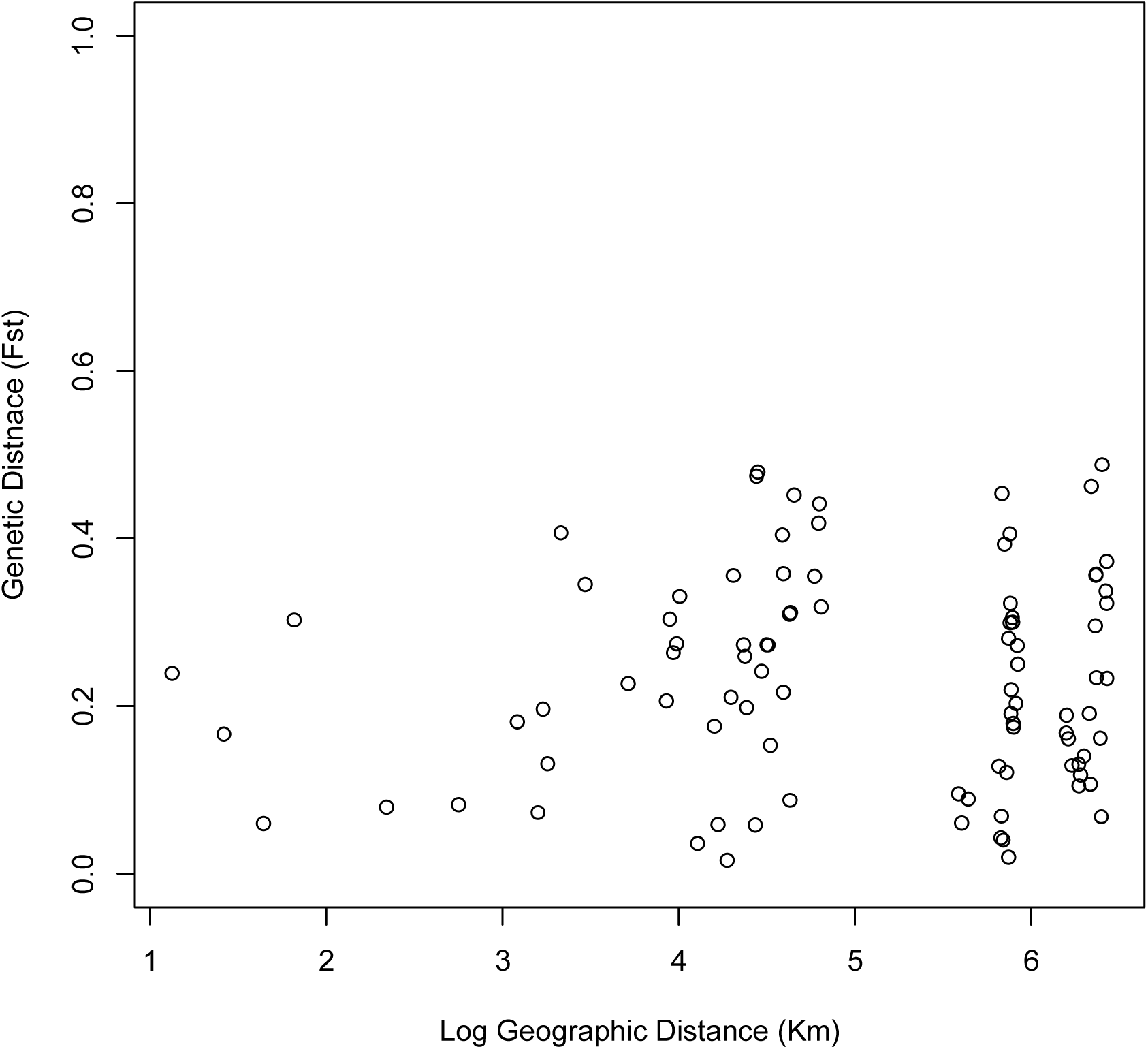
Pairwise genetic distance (*F_ST_*) of Bull Trout sampled within the Athabasca River basin plotted agains site’s the log pairwise geographic river distance (km)

## Discussion

### Overview of population structure of Bull Trout

Population genetics is an important conservation tool for understanding the genetic health of vulnerable populations (Tiffin & Ross-Ibarra 2014). In this study, Bull Trout exhibited patterns of high inter-population genetic differentiation within river basins with the majority of variation explained at the population level, a pattern found in other Salmonids in highly fragmented systems using a similar suite of markers (DeHaan *et al*. 2005; Ardren *et al*. 2011). All metrics of genetic differentiation and diversity that we considered support the conclusion that all river basins have similar levels of diversity, albeit with unique alleles and allele frequencies in each that differentiate the basins. The Athabasca river basin was found to contain several genetic clusters that did not correspond to the differences based on drainage of origin. These clusters were identified based on allele frequencies (by STRUCTURE), differentiation (*F_ST_, F_IS_*, and private alleles) and genetic diversity (*H_E_*and allelic richness). The majority of genetic variation was explained within each population with less variation explained among populations, and even less variation explained by the separate river basins.

### River basin scale differentiation and diversity

Despite the finding that Bull Trout in the Athabasca and North Saskatchewan River basins have similar levels of heterozygosity (*H_E_*), STRUCTURE results showed that the two river basins are highly differentiated from one another, exemplified by populations being clustered based on grouping fish to their origin in the Western Arctic or Saskatchewan-Nelson DU. In both river basins, high heterozygosity values and low *F*_IS_ were observed, alluding to large genetic differences between populations, which is common for Bull Trout systems (Costello *et al*. 2003). On average, the Saskatchewan River basin had higher allelic richness than the Athabasca River basin. The higher allelic richness may be a result of their closer proximity to the suspected glacial refugium (the Columbia refuge on the southern edge of the Cordilleran Ice sheet in the late Pleistocene; McPhail & Lindsey 1986), resulting in fewer subsequent founder effects during post-glacial dispersal than the further dispersed Athabasca River basin fish. Similarly, the North Saskatchewan river basin also had greater presence of private alleles, suggesting that each population is more differentiated in terms of novel genotypes, although this may be an artefact of fewer populations sampled in this area. Because fewer sites were sampled in the North Saskatchewan and Bow river basins compared to the Athabasca river basin, our values for genetic diversity and allelic richness may be underestimated in the former two basins due to limited localized sampling in those areas.

Anticipated differentiation for Bull Trout between the Athabasca and Saskatchewan watersheds was confirmed by STRUCTURE (Figure 3). This supports the Committee on the Status of Endangered Wildlife in Canada’s (COSEWIC 2012) designation of the two river basins as separate conservation units, the Western Arctic and Saskatchewan-Nelson, despite common genetic lineage. Genetic lineage is just one method used to support this designation, but it gives rationale for considering how these two different populations of Bull Trout will respond to management strategies, regional differences and climate change in unique ways. Previous studies using microsatellite loci have illuminated regional genetic differentiation in Bull Trout, corroborating the designation of these groupings (Spruell *et al*. 2003, Taylor *et al*. 1999). This type of differentiation on a large scale is common among fish (McPhail & Lindsey 1986) due to different refugia, extended isolation, and limited gene flow (Avise 2004). Because the groups have been separated for an extended period of time, evolutionary processes (e.g., genetic drift purging or fixing mutations in the two river basin’s populations) have influenced the genetic differentiation and divergence of Bull Trout in these two river basins.

### Sub-basin structure within the Athabasca river basin

Within the Athabasca River basin, additional sub-structuring was found. Genetic differences between populations were high but consistent with other Bull Trout studies (Warnock *et al*. 2010). Within the Athabasca River basin, differentiation among groups was best explained when using the five distinct clusters that were detected (Table 2) compared to either the tributary or larger regional area that the sampling site was located in. For each cluster, individuals were assigned to clusters with a high proportion of their allele frequencies matching the cluster with the exception of the McLeod-Moon cluster, which showed signs of admixture among individuals of different sampling site origin (specifically Moon Creek and McLeod River). Within the Athabasca River basin alone, these five groups likely constitute a mid-level within a hierarchical pattern of genetic diversity, with the majority of genetic variation explained within each of the five clusters with less variation among groupings and even less by river basins. Given that much genetic variation is explained in the population level and the McLeod-Moon cluster shows signs of two different genotypes within its membership, it is possible that further sub-structuring may exist within these groups.

### Isolation by Distance

Despite the evidence of sub-structuring in Bull Trout sampled the Athabasca River basin, no evidence for isolation by distance was found. This result suggests that the patterns of dispersal are weak, as expected, and that recent or continuing gene flow is present between our sites (Slatkin 1993). In lower, more homogenous stretches of the Athabasca river basin, another Salmonid, Arctic Grayling (*Thymallus arcticus*, Pallas 1776) exhibits moderate isolation-by-distance patterns, which is thought to be due to a mix of their large geographic ranges and population sizes (Reilly *et al*. 2014). Another Salmonid, Mountain Whitefish (*Prosopium williamsoni*, Girard 1856), also tends to exhibit low isolation-by-distance trend, although this is attributed to large population sizes and high levels of gene flow, which prevent differentiation (Whiteley *et al*. 2006).

### Implications

Given that Bull Trout populations are already declining, it is important that their genetic diversity is conserved across the range because this diversity may contain the local adaptation to help populations endure stochastic events (Rieman and McIntyre 1995), or it may be the evolutionary potential to be drawn from (Frankham 2005). Shrinking populations on the periphery of the range are especially at risk of extirpation as a result of isolation and habitat fragmentation (Rieman & Myers 1997). At present, many Bull Trout populations are listed as ‘data deficient’ with little insight into regionally specific differences from the highly studied areas from which management plans are drawn (COSEWIC 2012). Establishing and assessing baseline levels of genetic diversity allows for comparisons and detection of change in future stocks in addition to evaluation of adaptive management efforts (Epifanio *et al*. 2003). Uncovering areas that are genetically distinct from one another and determining the level of differentiation in local river basins can help refine management strategies to achieve local conservation goals. In Alberta, the genetic integrity of fish species and populations is determined based on the degree of hybridization, genetic similarity to original stock, and genetic distinction (Rodtka 2009). In this study, we provide baseline genetic information from which to track genetic changes observed in Bull Trout populations in the Athabasca River basin. Our results suggest little hybridization with Brook Trout at present in this region, as no hybrids or backcrosses were observed or genetically detected. With regard to genetic distinctiveness, we found genetic sub-structuring within the Athabasca River basin. High pairwise *F_ST_* values between sites indicate differentiation between groups (Table 3). However, high heterozygosity *(H_E_)* values and AMOVA results indicate that genetic variance largely resides within the population level. Thus, although there is differentiation between groups, there appears to be sufficient genetic diversity that can be drawn upon, acting as evolutionary potential to allow adaptation to changing habitat conditions and long-term persistence.

## Acknowledgements

This research was supported by funding and in-kind support from Parks Canada Agency and Alberta Environment and Parks and was supported by NSERC, QEII Scholarship, ACA, and ASPB Funds. John Post, Shelley Alexander and Ward Hughson provided project guidance and support. Peter Peller provided GIS support. Lindsay Dowbush and Traudi Golla assisted with field sampling.

