## Supplemental Table 1 for "Population genetics of Bull Trout (*Salvelinus confluentus*) in the Upper Athabasca river basin"

**Supplemental Table 1:** Microsatellites loci targeted to determine levels of genetic differentiation and population structure of Bull Trout in the Athabasca River basin. Optimized multiplexes are based on size range (bp) and annealing temperature (°C) for each locus. Expected heterozygosity ($H_E$) is averaged over 19 sampling sites at each allele. Expected heterozygosity ($H_E$) and Sources are given for each locus

| **Multiplex** | **Locus** | **Size range (bp)** | **Anneal Temp (°C)** | **N** | $H_E$ | **Source** |
|---|---|---|---|---|---|---|
| 1 | Omm1128 | 228-356 | 56.5 | 19 | 0.69 | (Rexroad *et al.* 2001) |
| | Sco105 | 164-204 | 56.5 | 13 | 0.55 | WDFW unpublished |
| 2 | Smm22 | 165-296 | 52 | 47 | 0.33 | (Crane *et al.* 2004) |
| | Sco102 | 148-290 | 52 | 8 | 0.27 | WDFW unpublished |
| | Sco215 | 207-328 | 52 | 8 | 0.23 | (Dehaan & Ardren 2005) |
| 3 | Sfo18 | 139-155 | 56 | 7 | 0.69 | (Popowich *et al.* 2011) |
| | Sco216 | 153-363 | 56 | 18 | 0.71 | (Popowich *et al.* 2011) |
| | Sco220 | 149-410 | 56 | 39 | 0.02 | (Dehaan & Ardren 2005) |
| 4 | Sco106 | 194-328 | 57.5 | 24 | 0.84 | WDFW unpublished |
| | Sco109 | 198-514 | 57.5 | 51 | 0.81 | WDFW unpublished |
