## Supplemental Table 2 for "Population genetics of Bull Trout (*Salvelinus confluentus*) in the Upper Athabasca river basin"

**Supplemental Table 2:** Loci in populations that are out of Hardy Weinberg Equilibrium (A) and presence of null alleles (B).

| A) Hardy- Weinberg Equilibrium | | B) Null Allele | |
|---|---|---|---|
| **Population** | **Locus** | **Population** | **Locus** |
| Athabasca River | Sco 109 | Athabasca River | Sco 220 |
| Berland River | Sco102 | Colt Creek | Sco 102 |
| Gregg River | Sco 102 | Colt Creek | Sfo 18 |
| Kerkeslin Lake | Sfo 18 | Colt Creek | Sco 215 |
| Kerkeslin Lake | Sco 220 | Elbow River | Sco 109 |
| Kerkeslin Lake | Smm 22 | Gregg River | Sco102 |
| Miette Lake | Sfo 18 | Kerkeslin Lake | Sfo 18 |
| Moon Creek | Sco 102 | Kerkeslin Lake | Sco 220 |
| Moon Creek | Sfo 18 | Little Elbow River | Smm22 |
| Osprey Lake | Sco 220 | Little Berland River | Sfo 18 |
| Colt Creek | Sco 102 | Little Berland River | Sco 220 |
| Colt Creek | Sfo 18 | McLeod River | Sco 215 |
| Elk Creek | Sfo 18 | Miette Lake | Sfo 18 |
| Elbow River | Sco 109 | Miette Lake | Sco 215 |
| Little Elbow River | Sco 102 | Moon Creek | Sco 102 |
| Little Elbow River | Smm 22 | Moon Creek | Sfo 18 |
| Storm Creek | Omm 1128 | Osprey Lake | Sco 220 |
| Storm Creek | Sco 109 | Storm Creek | Sco 109 |
