## Supplemental Table 3 for "Population genetics of Bull Trout (*Salvelinus confluentus*) in the Upper Athabasca river basin"

**Supplemental Table 3**: Expected heterozygosity, allelic richness and total private alleles at each sampling site. See population codes in abbreviations (Abr) in Table 1.

| Population Code | Expected Heterozygosity | Total Private Alleles | Allelic Richness | $F_{IS}$ Values |
|---|---|---|---|---|
| AR | 0.600 | 2 | 8.6 | 0.087 |
| BR | 0.534 | 1 | 6.1 | -0.106 |
| DC | - | - | - | - |
| GR | 0.473 | 2 | 4.4 | -0.027 |
| JL | 0.583 | 2 | 3.6 | 0.01 |
| KC | 0.589 | 4 | 8.4 | 0.044 |
| KL | 0.634 | 1 | 2.9 | 0.057 |
| LBR | 0.541 | 0 | 6.3 | 0.152 |
| MC | 0.716 | 1 | 5.4 | 0.047 |
| ML | 0.736 | 1 | 6.2 | -0.029 |
| MR | 0.483 | 1 | 6.6 | 0.094 |
| OL | 0.649 | 1 | 4.9 | -0.122 |
| RC | 0.530 | 0 | 2.8 | 0.076 |
| UL | 0.538 | 4 | 5.3 | 0.087 |
| CC | 0.69 | 0 | 8.1 | 0.143 |
| EC | 0.552 | 4 | 6.4 | 0.04 |
| ER | 0.571 | 0 | 3.7 | 0.07 |
| LER | 0.566 | 9 | 6.9 | -0.024 |
| PL | 0.468 | 8 | 7 | -0.044 |
| STC | 0.553 | 0 | 7.3 | 0.035 |
